# Endocytic accessory proteins assemble clathrin while simultaneously destabilizing protein condensates

**DOI:** 10.64898/2026.04.24.720715

**Authors:** Brandon Malady, Susovan Sarkar, Liping Wang, Zuzana Kadlecova, Eileen M. Lafer, David Owen, Jeanne C. Stachowiak

## Abstract

During endocytosis a dynamic protein network is responsible for assembly of the clathrin coat. Early in the process the Eps15 protein is thought to undergo liquid-liquid phase-separation to form biomolecular condensates. These condensates facilitate clathrin assembly, producing clathrin-coated vesicles that are ultimately excluded from the condensate as the vesicles depart from the plasma membrane. An array of distinct clathrin accessory proteins drive this process, yet their interactions with protein condensates have not been explored. Here we show that clathrin accessory proteins promote the assembly of the clathrin lattice by simultaneously destabilizing protein condensates. By adding diverse accessory proteins to Eps15 condensates, we observed that accessory proteins which cross-linked Eps15 proteins stabilized condensates, while accessory proteins that compete with Eps15-Eps15 interactions destabilized condensates. Interestingly, accessory proteins that destabilized condensates also promoted the assembly of clathrin and its exclusion from condensates. In contrast, accessory proteins that stabilized condensates opposed exclusion of clathrin. Finally, we demonstrated that decreasing condensate stability enhanced clathrin assembly and exclusion, even when no additional clathrin assembly motifs were introduced. Taken together, these results help to explain how condensates can catalyze the assembly of coated vesicles by providing both a substrate for their initiation and a driving force for their exclusion and ultimate departure.

**Statement of Significance:** This work is significant because it reveals how biomolecular condensates could actively drive, rather than merely host, clathrin-mediated endocytosis. By showing that clathrin accessory proteins can tune condensate stability to simultaneously promote clathrin lattice assembly and exclusion, this study uncovers a physical mechanism that could link phase separation to membrane trafficking. The finding that condensate destabilization alone enhances clathrin exclusion reframes condensates as dynamic regulators that could provide both an initiation platform and a force for vesicle maturation and release. More broadly, this work advances our understanding of how cells coordinate assembly and disassembly of complex macromolecular structures through regulated changes in condensate material properties.

## Introduction

Clathrin-mediated endocytosis (CME) is a major route for nutrient uptake, receptor recycling, and regulation of receptor signaling^1–3^. CME emerges from the cooperative assembly of a dynamic network of accessory proteins and clathrin triskelia at the plasma membrane. Recent work suggests that the early endocytic initiator protein, Eps15, undergoes liquid–liquid phase separation (LLPS) to form condensates that both concentrate the endocytic machinery and promote assembly of the clathrin lattice^4,5^, an observation that is consistent across mammalian^5^, yeast^6^, and plant cells^7^. Despite the increasing implication of biomolecular condensates in CME, the biophysical relationship between condensate properties and assembly of clathrin-coated vesicles remains poorly defined^5,8^. In particular, condensates must reconcile two seemingly opposing requirements of CME. They must be sufficiently cohesive to nucleate and organize the clathrin coat, yet they must remain fluid enough to allow rapid protein exchange, membrane deformation, and clathrin assembly. Precise tuning of condensate properties is likely required to meet these competing goals^9^.

Eps15 is thought to arrive early during CME, where it helps to initiate assembly of coated vesicles^10,11^. The domain structure of Eps15 includes three Eps15 homology (EH) domains at its N-terminus, which recognize NPF-containing ligands, followed by a central coiled-coil segment that mediates Eps15 dimerization, and finally a disordered C-terminal region, which contains multiple DPF motifs and other short linear motifs that engage other accessory proteins^11–13^**(SFig. 1)**. In isolation in vitro, Eps15 undergoes LLPS to form dynamic condensates through multivalent interactions with other Eps15 molecules^5^. Previous work has demonstrated that interactions between Eps15’s EH domains and its disordered C-terminus, along with coiled-coil mediated dimerization, are critical for assembly of Eps15 condensates. Interestingly, while soluble Eps15 does not drive clathrin assembly^4,14^, clathrin concentrates and assembles within Eps15 condensates^4^. The resulting clathrin assemblies are then dynamically excluded to the condensate periphery, such that clathrin and Eps15 spontaneously separate from one another *in vitro*.

In cells, Eps15 interacts with a diverse set of accessory proteins that can either reinforce or disrupt the multivalent Eps15–Eps15 interactions required for phase separation^5,8,15^. How might accessory proteins modify the stability of protein condensates and their ability to assemble clathrin? To address this question, we focused on accessory proteins that bound directly to Eps15 but differed in their potential for self-interaction. Proteins with a strong propensity for self-interaction, such as Fcho2 and Hrb, formed rigid, condensate-like structures under our *in vitro* conditions, such that they were able to increase the stability and decrease the fluidity of Eps15 condensates. In contrast, proteins with a weak propensity for self-interaction, such as AP2 and Epsin1, remained highly soluble, such that they decreased the stability and increased the fluidity of Eps15 condensates. Interestingly, accessory proteins that stabilized condensates inhibited the exclusion of clathrin lattices to the condensate surface, while accessory proteins that destabilized condensates promoted clathrin’s assembly and exclusion. These results suggest that the capacity of accessory proteins to drive assembly and departure of coated vesicles may depend on their ability to simultaneously disrupt protein condensates.

## Results

### Clathrin accessory proteins modify Eps15 condensate density

We began by forming condensates of Eps15 through the addition of 3% w/v polyethylene glycol (PEG 8000) to a solution containing 20 μM Eps15. Here PEG was used to mimic the crowded environment of the cytosol^16^. All condensates were formed in TNEET buffer, pH 7.5, 20 mM Tris-HCl, 150 mM NaCl, 1 mM EDTA, 1 mM EGTA, 5mM TCEP. We measured the fluorescence intensity of Eps15 in condensates relative to the dilute phase to determine the partitioning (Kp) of Eps15 within condensates, which is proportional to the relative protein density in the two phases (**Fig. 1A**). In the absence of additional accessory proteins, Eps15 had a Kp value of 31 ± 6.

**Figure 1.**
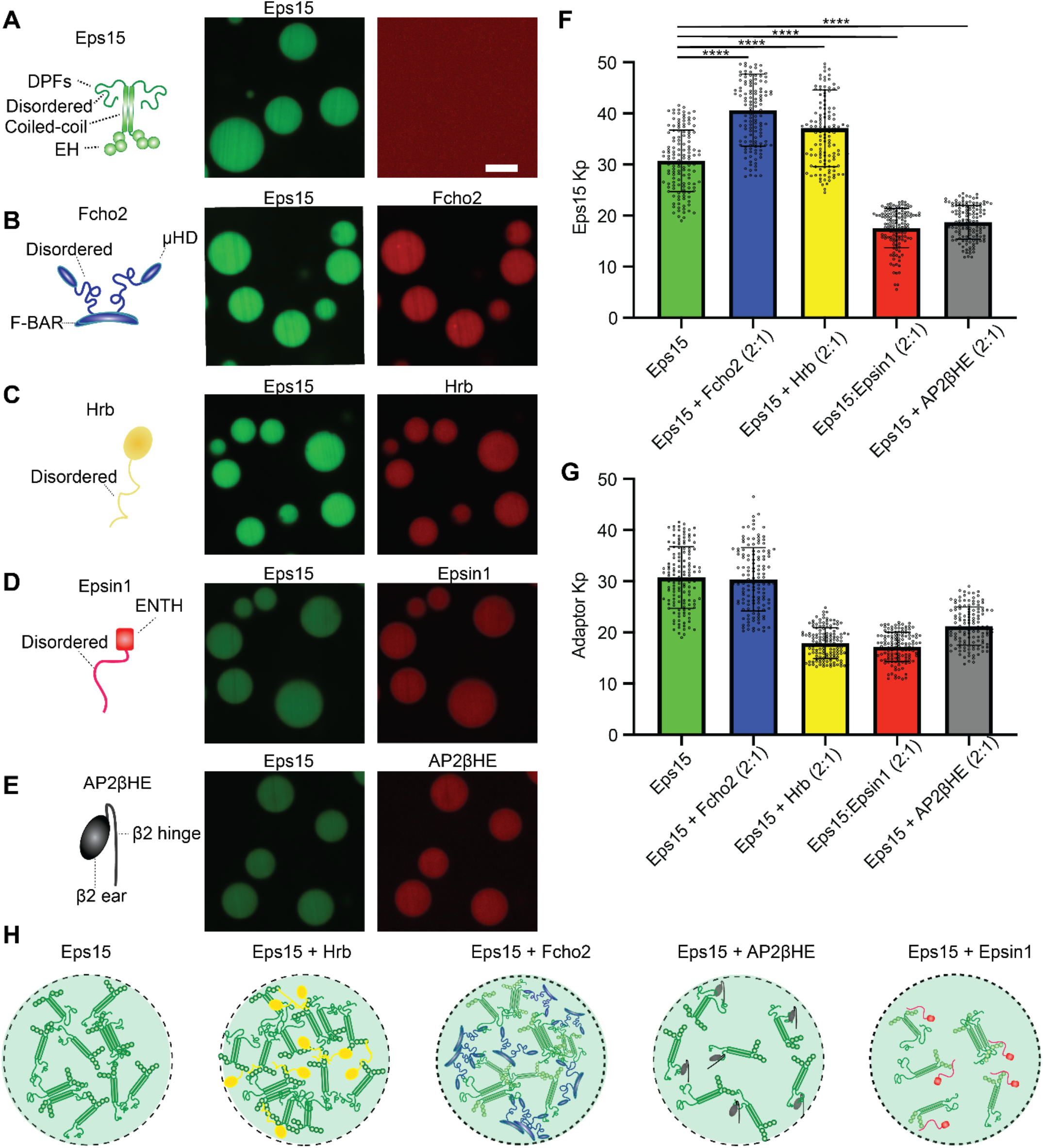
Clathrin accessory proteins modify Eps15 condensate density. **A-E)** Representative confocal microscopy images of Eps15 condensates with and without accessory proteins, n=3 biological replicates. For Eps15 only condensates in A) the red channel was included to demonstrate negligible fluorescence bleed through between channels. All condensates were formed in TNEET buffer (pH 7.5, 20 mM Tris-HCl, 150 mM NaCl, 1 mM EDTA, 1 mM EGTA, 5mM TCEP) with 20 μM Eps15 and 10 μM of accessory proteins when present. Scale bar is 5 micrometers. **F)** Bar Chart showing the partitioning (Kp) measurements for Eps15 into condensates with varying accessory proteins. **G)** Bar Chart showing the partitioning (Kp) measurements for accessory proteins into Eps15 condensates. For C) and D) 150 condensates were measured across n=3 separate biological replicates. **H)** Graphical depictions showing the effect of accessory proteins on the density of Eps15 within condensates. Welch’s two-tailed t-test was used to evaluate significance: ns denotes no significance, * denotes *p* ≤ 0.05, ** denotes *p* ≤ 0.01, *** denotes *p* ≤ 0.001 and **** denotes *p* ≤ 0.0001. GraphPad Prism Version 10.3.0 was used for statistical analysis.

In parallel, we examined the propensity for self-association of several distinct accessory proteins under these conditions: Fcho2, Hrb, AP2 (beta hinge and ear domains), and Epsin1 (**SFig 1**). Fcho2 is an early-arriving endocytic accessory protein that colocalizes with Eps15 at nascent endocytic sites^10,17^ and facilitates recruitment of downstream accessory proteins like AP2^18^. Its domain architecture consists of an N-terminal F-BAR domain that binds to membranes, a central disordered linker, and a C-terminal μ-homology domain^19^ (**SFig. 1**). The μ-homology domain of Fcho2 binds to DPF motifs in Eps15’s intrinsically disordered C-terminus^10,17^. Fcho2 forms dimers and higher order oligomers through its F-bar domain, creating the potential for multivalent interactions with Eps15 to establish an interconnected network^5,10,17^. In line with previous reports^5^, Fcho2 formed irregular, rigid condensates under our *in vitro* conditions, confirming a tendency toward self-interaction (**SFig. 2)**.

Hrb (HIV-1 Rev binding protein) is an accessory protein that colocalizes with Eps15 to promote cargo sorting^20,21^. Hrb consists of an N-terminal ArfGAP domain and a disordered C-terminal tail, which contains four NPF motifs, along with FG repeats throughout its sequence that may contribute to nuclear transport ^20–22^. Hrb interacts with Eps15 through its NPF motifs, which bind to Eps15’s EH domains^20^. Under our in vitro conditions, we found that Hrb formed irregular, rigid aggregates, similar to those observed with Fcho2 (**SFig. 2**).

Epsin1, an accessory protein in CME, binds ubiquitinated cargoes through ubiquitin interacting motifs while promoting membrane curvature essential for vesicle invagination^8,23^. Its architecture includes the epsin N-terminal homology (ENTH) domain which senses PI(4,5)P2 lipids and inserts an amphipathic helix into the plams membrane ^24^, followed by a C-terminal intrinsically disordered domain that contains tandem UIMs for ubiquitin recognition, DPW motifs for clathrin and AP2 recruitment, and three NPF motifs that bind to EH-domains in proteins such as Eps15^8,25,26^ (**SFig.1, SFig. 3**). In contrast to Fcho2 and Hrb, Epsin1 proteins are thought to repel one another, owing to the high net negative charge of Epsin1’s IDR domain and the lack of self-interaction domains within the protein^8,27^. In line with previous reports^8^, Epsin1 remained highly soluble under our *in vitro* conditions (**SFig. 2**).

Lastly, we investigated the self-interaction potential of the β-ear and hinge region of Adaptor protein complex 2 (AP2βHE)^4,28^. AP2 is a central endocytic accessory protein that links cargo and PI(4,5)P2 to the clathrin coat^29^. Its β2 hinge binds directly to clathrin to facilitate assembly of the clathrin lattice^28^. AP2βHE is known to bind to Eps15 via multiple FxDF motifs in Eps15’s C-terminal disordered region, which recognize a binding site on the side of the β2-appendage domain^28^. Owing to its lack of self-interacting motifs, we expected AP2βHE to remain highly soluble under our *in vitro* conditions (**SFig. 2**).

Having established their relative potential for self-interaction under our *in vitro* conditions, We asked how these accessory proteins might modulate the density of Eps15 proteins within condensates. We began with the accessory proteins that exhibited a strong propensity for self-interaction, Fcho2 and Hrb. Upon forming condensates composed of 20 μM Eps15 and 10 μM Fcho2, we found that Fcho2 was strongly recruited into condensates (Kp = 30 ± 6), resulting in an increase in the partitioning of Eps15 to the condensate phase (Kp = 41 ± 7) relative to condensates consisting of Eps15 alone (31 ± 6, **Fig. 1B**). Similarly, with condensates composed of 20 μM Eps15 and 10 μM Hrb, we observed strong Hrb partitioning (Kp = 17 ±3) and an increase in the relative partitioning of Eps15 within condensates (Kp = 37 ± 7) relative to Eps15-only condensates (**Fig. 1C**).

We next investigated how accessory proteins that lacked strong self-interactions, Epsin1 and AP2βHE, affected the density of Eps15 within condensates. We observed robust partitioning of Epsin1 within Eps15 condensates (Kp = 17 ± 3), and a corresponding decrease in Eps15 partitioning (Kp = 18 ± 4) for condensates formed with 20 μM Eps15 and 10 μM Epsin1, relative to Eps15-only condensates (**Fig. 1D**). This observation agrees with a previous study in which Epsin1 was found to destabilize Eps15 condensates^8^. Similarly, in condensates composed of 20 μM Eps15 and 10 μM AP2βHE, AP2βHE displayed robust partitioning within condensates (Kp = 21± 4), resulting in a decrease in Eps15 partitioning within condensates (Kp = 19 ± 3) relative to Eps15-only condensates (**Fig. 1E**).

The changes in relative partitioning of Eps15 within condensates upon addition of specific accessory proteins revealed that self-interacting accessory proteins (Fcho, Hrb) tended to increase Eps15 network density, presumably by cross-linking Eps15 dimers, while accessory proteins that lacked strong self-interactions (AP2βHE, Epsin1) decreased Eps15 network density, presumably by competing with Eps15-Eps15 interactions (**Fig. 1F-H**)^4,5,10^. Specifically, while all four accessory proteins have the potential to interact with Eps15 in ways that could, in principle, compete with Eps15-Eps15 interactions, those with strong self-interactions are more likely to present a multivalent scaffold for binding of multiple Eps15 proteins, resulting in stabilization of condensates. In contrast, those that lack strong self-interactions have the potential to compete with Eps15-Eps15 interactions in a way that disrupts condensates by increasing the partitioning of Eps15 dimers in solution. We would expect these changes in the connectivity of Eps15 proteins within condensates to impact condensate fluidity. Therefore, we next investigated the impact of accessory proteins on the liquid-like properties of Eps15 condensates.

### Recruitment of accessory proteins alters condensate physical properties

Previous work suggests that Eps15 condensates are liquid-like, as they fuse and re-round rapidly upon contact, and that the proteins within them undergo rapid exchange, as evidenced by rapid recovery after photo-bleaching^5,30^. We evaluated the impact of accessory proteins on these dynamic properties. We formed condensates of Eps15 in the absence and presence of the accessory proteins tested in Figure 1. We measured the fusion dynamics of condensates in image series acquired at one frame per second, restricting our analysis to fusion events between condensates of 3-5 μm diameter for all compositions in **Figure 2**. Condensates composed solely of 20 μM Eps15 merged and fully re-rounded within 2.6 ± 0.5 seconds on average.

**Figure 2.**
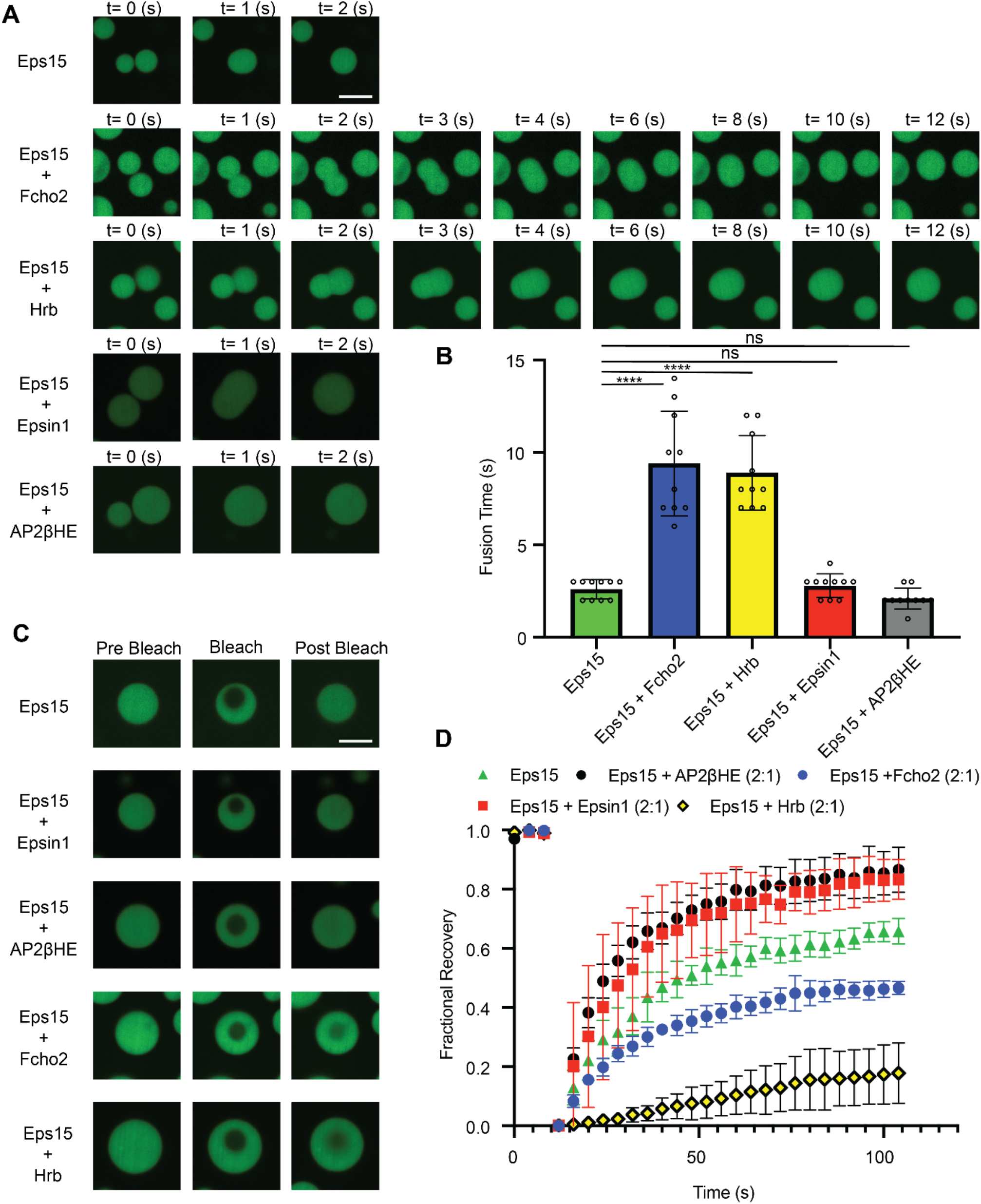
Recruitment of accessory proteins alters condensate physical properties. **A)** Representative confocal microscopy images showing merging and re-rounding of Eps15 condensates with and without additional accessory proteins, n=3 separate biological replicates for each condition. **B)** Bar chart depicting merging and re-rounding times for each condensate system. For each condition, 10 fusion events were monitored across 3 separate biological replicates. Error bars are standard deviation. **C)** Representative confocal microscopy images depicting fluorescence recovery after photobleaching events for each composition. Images show the pre-bleach, bleach, and post bleach timepoints at 2 minutes after photobleaching. **D)** Curves depicting the fluorescence recovery after photobleaching for condensates of the indicated compositions. Recovery curves represent the average recovery across 3 separate replicates. Error bars are standard deviation. All condensates were formed in TNEET buffer (pH 7.5, 20 mM Tris-HCl, 150 mM NaCl, 1 mM EDTA, 1 mM EGTA, 5mM TCEP) with 20 μM Eps15 and 10 μM of accessory proteins when present. For statistics, Welch’s two-tailed t-test was used: ns denotes no significance, * denotes *p* ≤ 0.05, ** denotes *p* ≤ 0.01, *** denotes *p* ≤ 0.001 and **** denotes *p* ≤ 0.0001. GraphPad Prism Version 10.3.0 was used for statistical analysis.

We next examined the impact of accessory proteins with strong self-interactions. Condensates composed of 20 μM Eps15 + 10 μM Fcho2 or 20 μM Eps15 + 10 μM Hrb merged and re-rounded more slowly, within 8.9 ± 2 seconds and 9.4 ± 2.8 seconds, respectively (**Fig. 2A**). This result is in line with the ability of self-interacting accessory proteins to increase condensate stability as observed above. In contrast, addition of accessory proteins that lacked strong self-interactions, Epsin1 and AP2βHE, merged and re-rounded over similar timescales to condensates consisting of Eps15 alone, 2.8 ± 0.6 seconds and 2.1 ± 0.6 seconds, respectively (**Fig. 2B**).

As an orthogonal measure of their fluidity, condensates were photobleached in regions with an approximate area of 3 μm in their center and their fluorescence recovery over 2 minutes was monitored (**Fig. 2C**). Condensates composed of 20 μM Eps15 alone recovered by 66 ± 4% with a half-life (τ1/2) of 17 seconds (**SFig. 4**). In contrast, condensates composed of 20 μM Eps15 and 10 μM Fcho2 recovered by 47 ± 2% with τ1/2 of 18 seconds, while condensates composed of 20 μM Eps15 and 10 μM Hrb recovered by 18 ± 1% and had a substantially longer τ1/2 of 45 seconds (**SFig. 4**). Reduced recovery in the presence of Fcho2 and Hrb is consistent with the ability of these accessory proteins to cross-link the Eps15 network. Additionally, we investigated the effect of the non-self-interacting accessory proteins, Epsin1 and AP2βHE, in FRAP experiments. Condensates composed of 20 μM Eps15 and 10 μM Epsin1 recovered by 87 ± 7% with a τ1/2 of 14 seconds, indicative of increased liquidity relative to Eps15-only condensates. Similarly, condensates composed of 20 μM Eps15 and 10 μM AP2βHE recovered by 86 ± 8% with a τ1/2 of 11 seconds. (**Fig. 2D, SFig. 4**).

Taken together, these data suggest that self-interacting accessory proteins reduce the fluidity and molecular exchange within condensates, likely through increased crosslinking of Eps15 proteins, while accessory proteins that lack strong self-interactions increase condensate fluidity and molecular exchange, likely by competing with Eps15-Eps15 interactions. Building on these results, we next investigated the impact of accessory proteins on interactions between condensates and clathrin.

### Accessory proteins tune clathrin assembly and exclusion from condensates

We previously demonstrated that Eps15 condensates facilitate the assembly of clathrin triskelia, which initially partition to the condensate interior, where they assemble into lattices that are gradually excluded to the condensate surface^4^. This peripherally excluded clathrin failed to recovery after photobleaching, indicating slow molecular exchange, as would be expected for a stably assembled lattice. Additionally, electron microscopy revealed mature clathrin assemblies and lattices on the surfaces of Eps15 condensates^4^. Building on these findings, we sought to determine the impact of accessory proteins on the ability of condensates to assemble and exclude the clathrin lattice. Toward this goal, we first tested each accessory protein’s native ability to assemble clathrin under dilute conditions, in the absence of protein condensates. Here we employed dynamic light scattering, which is known to show a significant shift toward larger hydrodynamic radii upon clathrin assembly^4,14^. Using using pH to control the assembly state of clathrin^4,29^ we confirmed that clathrin triskelia, which are stably disassembled at pH 7.5, have a hydrodynamic diameter of 39 ± 1 nm and clathrin baskets, which assemble stably at pH 6.8, have a hydrodynamic diameter of 120 ± 3 nm at a clathrin concentration of 200 nM, with a 1:1 ratio of clathrin heavy chain (CHC) and and clathrin light chain (CLC), in line with our previous findings^4^.

We also determined the hydrodynamic diameter of each accessory protein alone in solution at a fixed concentration of 10 μM. Here 3% w/v PEG was absent so that condensates and aggregates did not form, and all accessory proteins remained in solution. Eps15, Epsin1, AP2βHE, Fcho2, and Hrb had average diameters of 29 ± 1 nm, 16 ± 2 nm, 15 ± 1 nm, 23 ± 1 nm, and 13 ± 3 nm respectively (**Fig. 3A**), all well below that of clathrin baskets. We then prepared samples of 10 μM of each accessory protein with 200 nM clathrin, the same protein concentrations used in the condensate assays. We found that Epsin1 and AP2βHE led to the assembly of particles with diameters similar to clathrin baskets, 98 ± 28 nm and 86 ± 4 nm, respectively. These results are consistent with the known ability of these accessory proteins to facilitate clathrin assembly^25,28,31^. In contrast DLS measurements of the hydrodynamic radii for 10 μM Hrb or 10 μM Fcho2 with 200 nM of clathrin triskelia had average hydrodynamic radii of 41 ± 2 nm and 34 ± 1 nm. These values are larger than the adaptor proteins alone, but smaller than clathrin baskets. They are similar to the measured hydrodynamic radii of individual clathrin triskelia (39 ± 1 nm), which, despite being a minority component in the solution, are likely to dominate the dynamic light scattering data, given the strong dependence of light scattering on particle size^32^. Taken together, these data confirm the known ability of Epsin1 and AP2βHE to drive clathrin assembly in the absence of protein condensates, while suggesting that Fcho2 and Hrb lack this ability.

**Figure 3.**
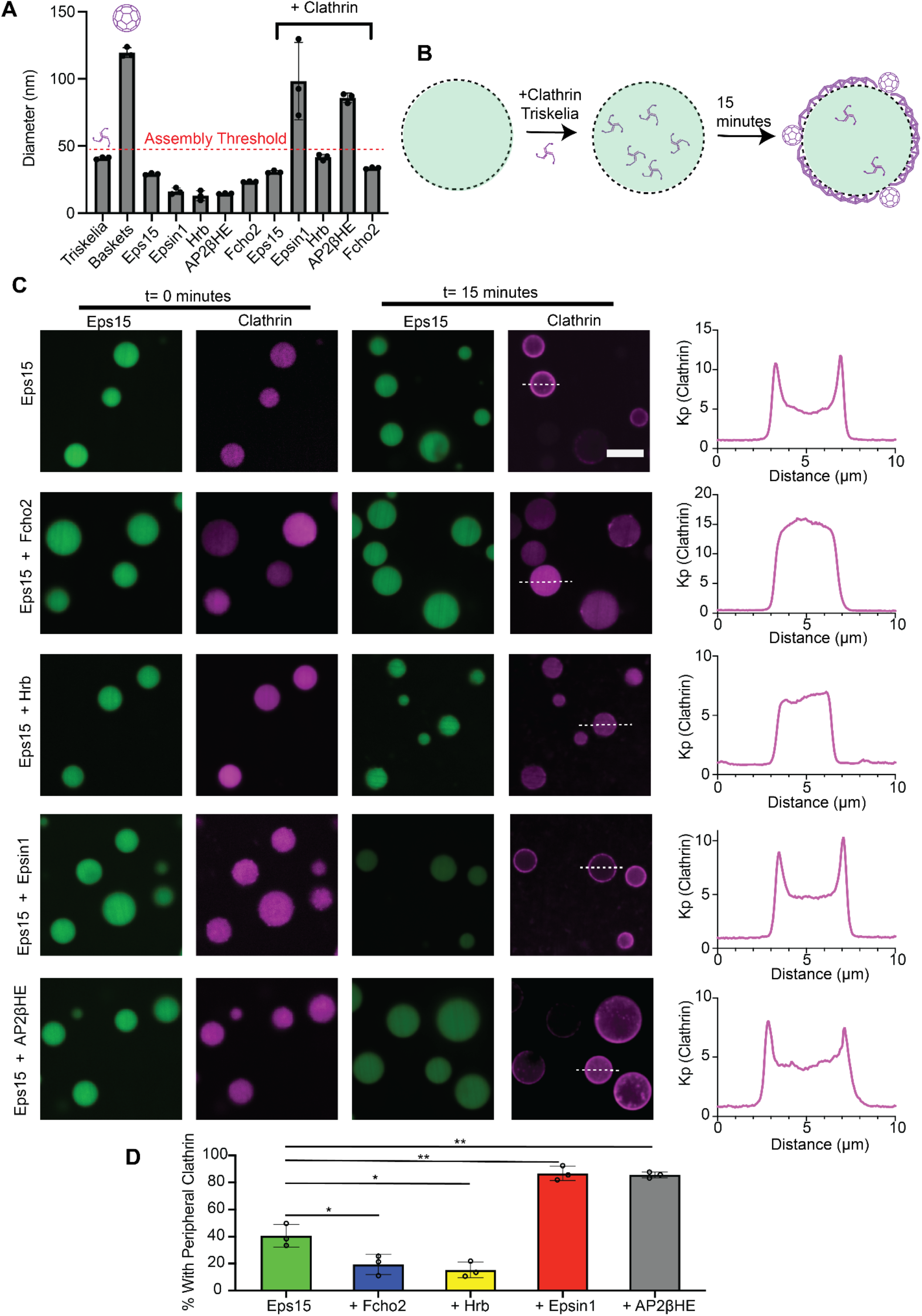
Accessory proteins tune clathrin assembly in Eps15 condensates. **A)** Dynamic light scattering data depicting the average hydrodynamic diameter ± s.d. for each of the indicated conditions. All conditions were measured in pH 7.5 TNEET buffer except for the clathrin baskets which were formed with pH 6.8 TNEET, n = 3 independent replicates with 15 measurements averaged per replicate. For DLS protein concentrations were 10 μM of the corresponding accessory protein along with 200nM of clathrin triskelia when present. **B)** Graphical depiction showing condensate-mediated clathrin assembly and experimental scheme for C) and D) along with representative confocal microscopy images showing the initial partitioning within and subsequent exclusion of clathrin from condensates. **C)** Representative confocal microscopy images depicting the initial accumulation of clathrin within endocytic condensates and subsequent formation of clathrin around endocytic condensates of various network compositions 15 minutes following clathrin addition along with corresponding line profiles, n=3 separate biological replicates. The scale bar is 5 micrometers. **D)** Quantification of peripheral clathrin accumulation assay outlined in B and C. Condensates are considered to have clathrin excluded to the periphery if the signal intensity at the periphery was 1.5x that of the interior, n=3 biological replicates, bar charts depict the average percentage of condensates with clathrin excluded to the periphery at each replicate ± s.d. All condensate experiments were conducted with 20 μM Eps15 + 10 μM of the corresponding accessory protein and 200 nM of clathrin triskelia. For statistics Welch’s two-tailed t-test was used: ns denotes no significance, * denotes *p* ≤ 0.05, ** denotes *p* ≤ 0.01, *** denotes *p* ≤ 0.001 and **** denotes *p* ≤ 0.0001. GraphPad Prism Version 10.3.0 was used for statistical analysis.

We next sought to determine how the presence of each accessory protein impacted the ability of Eps15 condensates to drive clathrin assembly and subsequent exclusion from the condensate network. For this purpose, we formed condensates consisting of accessory proteins as described above. Condensates were added to a passivated coverslip and allowed to settle for 10 minutes before clathrin, labeled with Alexa 647 NHS ester, was added to achieve a final clathrin concentration of 200 nM. Upon its addition, clathrin initially partitioned uniformly to condensates, followed by gradual repartitioning to the condensate periphery (**Fig. 3B,C**), as observed previously^4^. We collected confocal microscopy images at a fixed time point of 15 minutes following clathrin addition and quantified the fraction of condensates with peripheral accumulation of clathrin (**Fig. 3C**). Condensates were determined to have peripheral clathrin if the fluorescence intensity of clathrin around the condensate periphery was greater than 1.5x that within condensates (**Fig. 3C**). Condensates formed by Eps15 alone had peripheral clathrin 41 ± 8% of the time. In contrast, condensates composed of 20 μM Eps15 and 10 μM Fcho2 or Hrb, accessory proteins with strong self-interactions, had peripheral clathrin 19 ± 7% and 15 ± 6% of the time, respectively, indicating that these accessory proteins disfavored peripheral accumulation of clathrin. In contrast, condensates composed of 20 μM Eps15 and accessory proteins with weak self-interactions and a native ability to assembly clathrin (10 μM of Epsin1 or AP2βHE) displayed peripheral accumulation of clathrin 87 ± 5% and 86 ± 2% of the time, respectively, a substantial increase relative to condensates composed of Eps15 alone (41 ± 8%). Together these observations suggest that condensates formed by the inclusion of self-interacting accessory proteins were less able to exclude clathrin assemblies, perhaps due to their reduced fluidity. In contrast, the more fluid condensates formed through the inclusion of non-self-interacting accessory proteins tended to promote exclusion of clathrin.

### Molecular exchange within condensates of accessory proteins and clathirn

To probe the ability of proteins to exchange within condensates composed of accessory proteins and clathrin, we performed FRAP experiments, which utilized the same condensate compositions and buffer conditions as the experiments in Figure 3. In condensates composed of Eps15 and self-interacting accessory proteins, Hrb or Fcho2, we showed above that clathrin remained predominantly within condensates, rather than being excluded to the condensate periphery (**Fig. 3C, 4A**). FRAP experiments revealed minimal molecular exchange of the clathrin within these condensates (**Fig. 4A-B**). In contrast, the Eps15 proteins within these condensates maintained some molecular exchange, albeit at lower levels (8% ± 5% or 7% ± 4% for condensates containing Hrb or Fcho2, respectively), compared to condensates consisting only of Eps15 and clathrin (38% ± 18%), (**Fig. 4A,C**). Notably, Eps15 exchanged even more completely (66 ± 4%) in the absence of clathrin, (**Fig 2C,D**). In line with our previous work, the reduction in Eps15 exchange upon addition of clathrin suggests that clathrin is assembling within condensates and thereby stabilizing them^4^.

**Figure 4.**
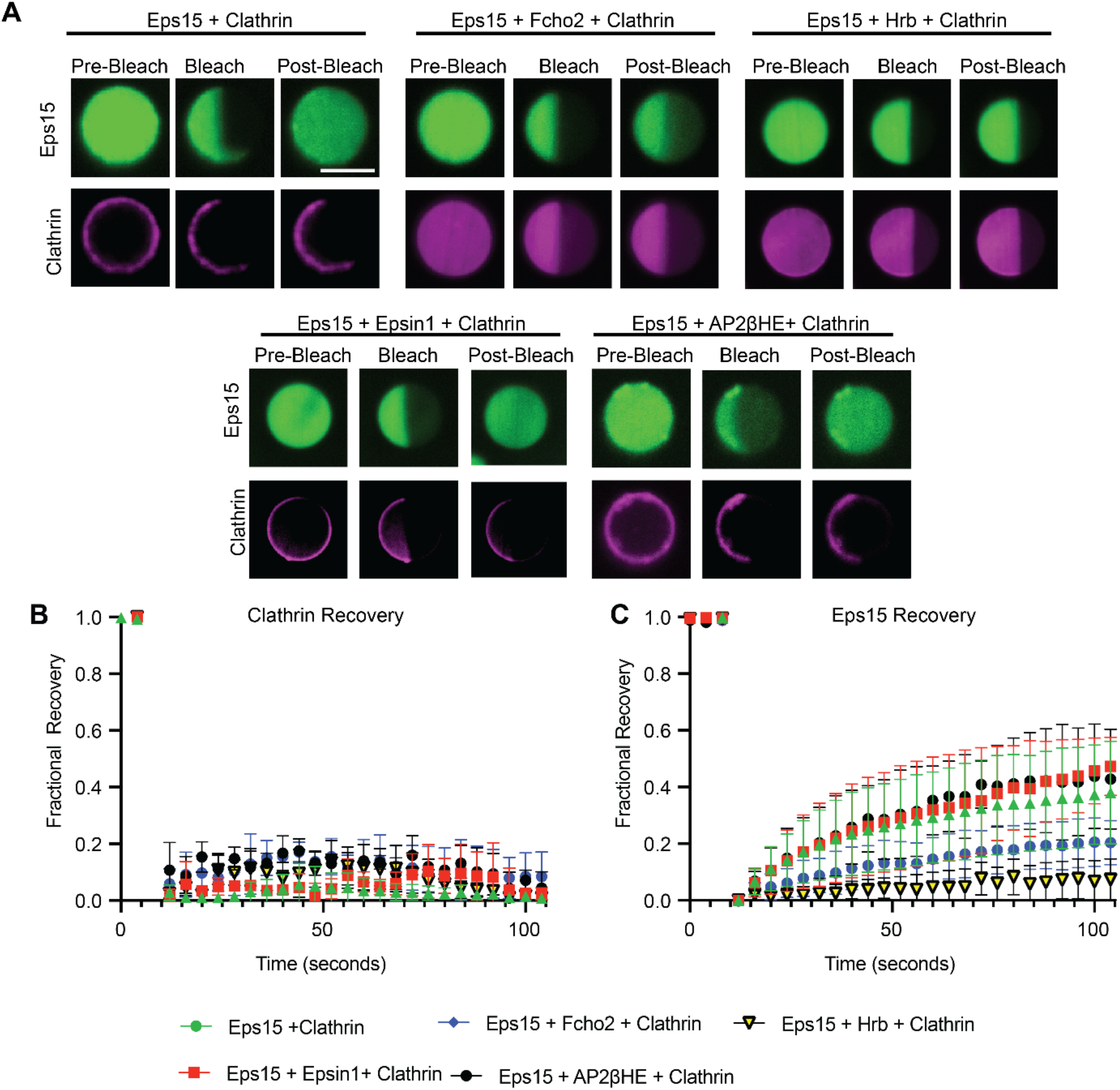
Molecular exchange within condensates of accessory proteins and clathirn. **A)** Confocal microscopy images showing the pre-bleach, bleach, and 2 minutes post-bleach frames for condensates consisting of adaptor proteins and clathrin. All condensates were formed in TNEET buffer pH 7.5 20 mM Tris-HCl, 150 mM NaCl, 1 mM EDTA, 1 mM EGTA, 5mM TCEP) with 20 μM Eps15 + 10 μM of the corresponding accessory protein and 200 nM of clathrin triskelia. Here contrast was adjusted to show the recovery post-bleaching, resulting in some of the pre-bleach images appearing saturated. **B)** Normalized recovery profiles for FRAP experiments for the clathrin channel for the condensate compositions in A). **C)** Normalized recovery profiles for FRAP experiments for the Eps15 channel for the network compositions in A). Scale bar is 5 μm. Recovery curves represent the average recovery across 3 replicates. Error bars are standard deviation.

In contrast, for condensates composed of Eps15 and non-self-interacting accessory proteins, Epsin1 or AP2βHE, we showed above that clathrin was substantially excluded to the condensate periphery (**Fig, 3C, 4A**). Here FRAP experiments revealed minimal molecular exchange of peripheral clathrin, suggesting that the excluded clathrin is stably assembled, in agreement with our previous findings, which correlated FRAP data with electron microscopy^4^. In contrast, Eps15 proteins within these condensate maintained the ability to exchange at substantial levels (47% ± 10% or 43% ± 17% for condensates containing Epsin1 or AP2βHE, respectively), an increase compared to condensates consisting only of Eps15 and clathrin (38% ± 18%), though still somewhat less than in condensates composed of only Eps15 (66 ± 4%) (**Fig. 4A,C**).

Taken together, the results in Figures 4 suggest that clathrin assembles to some degree within condensates composed of Eps15 and each of the four accessory proteins we have examined. This conclusion is supported by the lack of clathrin recovery after photo-bleaching, whether it was excluded to the condensate periphery, upon addition of Epsin1 or AP2βHE, or remained inside the condensate, upon addition of Hrb or Fcho2 (**Fig. 4B**). Further evidence comes from the reduction in Eps15 recovery for all condensate compositions upon addition of clathrin (**Fig. 4C**). Notably, addition of either Epsin1 or AP2βHE resulted in both (i) a substantial increase in exclusion of clathrin to the condensate periphery and (ii) increased molecular exchange of Eps15 in the presence of clathrin relative to condensates composed only of Eps15 and clathrin, in sharp contrast to addition of Hrb or Fcho2, which resulted in a substantial loss of Eps15 exchange. The correlation between clathrin exclusion and Eps15 exchange suggests that the ability of condensates to exclude clathrin assemblies likely requires that condensates remain highly fluid. However, the inherent ability of Epsin1 and AP2βHE to assemble clathrin makes it difficult to determine the contribution of increased condensate fluidity as opposed to specific interactions with these clathrin assembly proteins. Therefore, we next sought to isolate the contribution of increased condensate fluidity to the exclusion of clathrin assemblies from condensates.

### Increasing condensate fluidity enhances clathrin assembly and dynamic exclusion

We next sought to decouple the contributions of condensate fluidity and accessory protein content to the exclusion of assembled clathrin to the condensate periphery. For this purpose, we employed an Eps15 mutant which we expected to fluidize Eps15 condensates without increasing their biochemical capacity to assemble triskelia into lattices. Specifically, we added a version of Eps15 lacking the coiled-coil domain responsible for Eps15-dimerization (Eps15ΔCC) to condensates. Our previous work demonstrated that this mutant lacks the ability to assemble into protein condensates^5^. We expected that addition of Eps15ΔCC to condensates composed of wild type Eps15 would increase their fluidity by competing with interactions between dimers of wild type Eps15 (**Fig. 5A**). To evaluate this idea, we formed condensates composed of 20 μM Eps15 and 10 μM Eps15ΔCC. We observed Eps15ΔCC partitioning within condensates (Kp= 16 ± 3) and a corresponding decrease in partitioning of wild type Eps15 into condensates (Kp= 20 ± 8) relative to condensates consisting of wild type Eps15 alone (Kp= 31 ± 6), confirming the expected reduction in the density of wild type Eps15 within condensates. In addition, we tested a higher concentration of Eps15ΔCC relative to Eps15 by forming condensates composed of 20 μM Eps15 and 20 μM Eps15ΔCC. This 1:1 molar-ratio of Eps15 to Eps15ΔCC led to an even greater reduction in wild type Eps15 partitioning (Kp= 14.28 ± 2) within condensates along with increased Eps15ΔCC partitioning (Kp= 27 ± 4) (**Fig. 5B,C**). We then tested the liquidity of condensates formed with 20 μM wild type Eps15 and 10 μM Eps15ΔCC and found they displayed fast fusion times, merging and re-rounding within 2 seconds (**Fig. 5D**), faster than condensates composed of wild type Eps15 (2.6 seconds, **Fig. 2A,B**). We also used FRAP as described in Figure 2 to further assess the liquidity of condensates formed from 20 μM wild type Eps15 and 10 μM Eps15ΔCC (**Fig. 5E**). Here we observed an increase in fluorescence recovery after photobleaching for condensates with Eps15ΔCC (81 ± 6%) relative to condensates composed of wild type Eps15 alone (66 ± 4%) (**Fig. 5F**).

**Figure 5.**
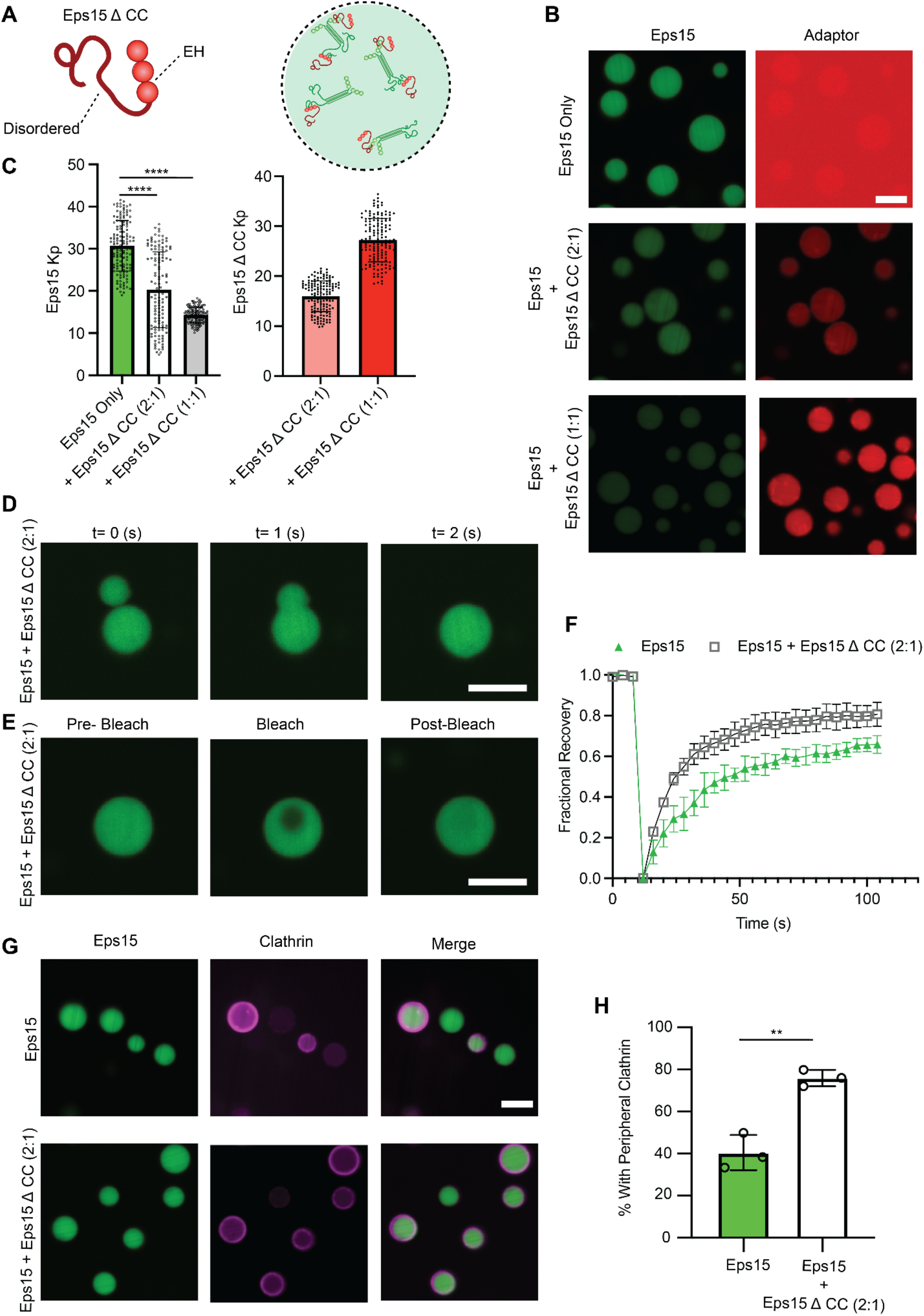
Increasing condensate fluidity enhances clathrin assembly and dynamic exclusion. A) Graphical depiction of Eps15 ΔCC mutant in which the coiled-coil dimerization domain is removed along with its graphical impact on network density. B) Representative confocal microscopy images showing the partitioning of Eps15 ΔCC into condensates as well as the effect on Eps15 auto partitioning. **C)** Quantification of Eps15 auto partitioning into condensates as a function of Eps15 ΔCC titration along with quantification of partitioning of Eps15 ΔCC into condensates. Here condensates were formed at 20 μM Eps15 along with either 10 μM Eps15 ΔCC or 20 μM Eps15 ΔCC as indicated. Bar charts represent average Kp ± s.d. For 150 measured condensates across n=3 separate replicates for each condition. **D)** Representative microscopy images depicting fast fusion of condensates formed with 20 μM Eps15 and 10 μM Eps15 ΔCC. **E)** Representative microscopy images depicting the recovery after photobleaching for condensates formed with 20 μM Eps15 and 10 μM of Eps15 ΔCC. **F)** Curves depicting the fluorescence recovery after photobleaching for condensates formed at the indicated conditions. Recovery curves represent the average recovery across 3 separate replicates. Error bars are standard deviations. All condensates were formed in TNEET buffer (pH 7.5 20 mM Tris-HCl, 150 mM NaCl, 1 mM EDTA, 1 mM EGTA, 5mM TCEP, 3% PEG) with 20 μM Eps15 and 10 μM of Eps15 ΔCC when present. **G)** Representative microscopy images of peripheral clathrin quantification assay as in figure 3 for Eps15 alone and Eps15 + Eps15 ΔCC. **H)** Quantification of clathrin exclusion assay outlined in figure 3. Condensates are considered to have clathrin excluded to the periphery if the signal intensity at the periphery was 1.5x that of the interior, n=3 biological replicates, bar charts depict the average percentage of condensates with peripheral clathrin at each replicate ± s.d. Experiments for the assay quantifying the amount of peripherally-excluded clathrin were conducted with 20 μM Eps15 + 10 μM of the corresponding accessory protein and 200 nM of clathrin triskelia. For statistics Welch’s two-tailed t-test was used: ns denotes no significance, * denotes *p* ≤ 0.05, ** denotes *p* ≤ 0.01, *** denotes *p* ≤ 0.001 and **** denotes *p* ≤ 0.0001. GraphPad Prism Version 10.3.0 was used for statistical analysis.

We next sought to determine to what extent enhanced condensate fluidity impacted the efficiency of condensate-mediated clathrin assembly. For this purpose, we returned to our clathrin assembly assay in Figure 3B. Here we added 200 nM of clathrin triskelia to condensates formed with 20 μM wild type Eps15 and 10 μM Eps15ΔCC (**Fig. 5G**). We observed an increase in the fraction of condensates with peripheral clathrin (76 ± 4%) compared to condensates composed of wild type Eps15 alone (41 ± 8%), an increase which approached that achieved through addition of Epsin1 or AP2βHE, which are known to contain strong clathrin assembly motifs (**Fig. 5H vs. Fig. 5D**). Taken together these results suggest that increasing condensate fluidity is itself a strong enhancer of clathrin assembly, which is orthogonal to the addition of strong clathrin assembly motifs.

## Discussion

It is increasingly clear that protein condensates composed of the mammalian accessory protein, Eps15, and its homologs in yeast and plants, play an important role in the early initiation of clathrin-mediated endocytosis^5–7^. Here we asked how clathrin’s diverse array of accessory proteins modify the stability and physical properties of Eps15 condensates, and ultimately how they impact the assembly and exclusion of clathrin lattices in the condensate environment. We show that accessory proteins such as Fcho2 and Hrb, which have strong self-interactions, tend to stabilize condensation of Eps15, likely through their capacity to bind simultaneously to multiple Eps15 proteins^10,20^. In contrast, we find that accessory proteins such as Epsin1 and AP2βHE, which have minimal self-interactions, tend to destabilize condensation of Eps15, likely by competing with Eps15’s self-interactions. Importantly, each of these proteins partitioned strongly into Eps15 condensates, underscoring Eps15’s potential role as a recruitment hub for the endocytic machinery^10^. The dichotomy between accessory proteins with and without strong self-interactions points to a framework in which crosslinking of Eps15 by proteins with strong self-association could build a scaffold that concentrates the endocytic machinery, which is ultimately disassembled by accessory proteins with weak self-association, once it has served its purpose. Specifically, Fcho2 and Hrb enhance Eps15 network density while decreasing network fluidity in our experiments. These behaviors likely arise from multivalent interactions with Eps15 such as binding between the μ-homology domains of the Fcho2 dimer with proline-rich stretch to nucleate CME^10,17^. Fcho2’s dimeric μ-homology domains^17^ and Hrb’s multiple NPF motifs^20^ could bridge multiple Eps15 molecules, boosting Eps15 partitioning and yielding slow-fusing condensates with suppressed molecular exchange, indicating increasingly stable protein networks. These properties align with the role of Fcho2 in stabilizing nascent endocytic sites^10^.

In contrast, accessory proteins with a low propensity for self-interaction, AP2βHE and Epsin1, reduce the density of the Eps15 network yielding droplets that merge more rapidly and display greater molecular exchange than droplets composed of Eps15 alone. These behaviors likely arise from the ability of these proteins to resist condensation while also competing with Eps15’s homotypic interactions^5,13^. Further, condensates that contained these accessory proteins showed a greater propensity to assemble and exclude clathrin lattices. However, it was unclear whether this capacity arose from the ability of AP2βHE and Epsin1 to fluidize Eps15 condensates versus their inherent ability to assemble clathrin lattices. Therefore, to isolate the contribution of condensate fluidity, we utilized a monomeric version of Eps15 (Eps15ΔCC) that competed with interactions between Eps15 dimers but lacked clathrin-assembly motifs. Addition of Eps15ΔCC to droplets composed of wild-type Eps15 resulted in a reduction in the density of wild-type Eps15 within condensates, which was accompanied by an increase in condensate fluidity, as evidenced by faster condensate fusion and higher rates of molecular exchange in comparison to condensates composed of wild-type Eps15 alone. Strikingly, this increased fluidity, without introducing additional clathrin assembly motifs, substantially increased the fraction of condensates that assembled and excluded clathrin, similar to the addition of AP2βHE or Epsin1. These results reveal that the ability of accessory proteins to enhance condensate fluidity is itself a predictor of the efficiency of clathrin assembly and exclusion. These findings suggest that optimal conditions for clathrin assembly and exclusion arise within protein networks that are cohesive enough to concentrate clathrin triskelia yet fluid enough to support their rapid rearrangement into lattices that are subsequently excluded from the condensate. More broadly, this framework suggests that cells could tune the kinetics and morphology of clathrin-coated structures by modulating accessory protein stoichiometry.

While our experiments were conducted *in vitro* without PI(4,5)P2 containing membranes, it is worth noting that binding of Fcho2 to PI(4,5)P2 is critical for localization of the Eps15 network to the plasma membrane^10,18^ and subsequent recruitment of downstream accessory proteins and clathrin. Furthermore, our experiments do not include endocytic cargo molecules, which will strongly influence the recruitment of specific accessory proteins. In particular, the presence of ubiquitinated cargo could shift the role of Epsin1 from disrupting to stabilizing Eps115 condensates^8^. Future work on how cargo molecules, membrane interactions, and additional accessory proteins modulate the properties of endocytic condensates will be important to develop a mechanistic understanding of clathrin-mediated endocytosis.

## Experimental Methods

### Plasmids

The plasmid encoding full-length Homo sapiens Eps15 (pET28a-6xHis-Eps15-FL) was generously provided by Tom Kirchhausen. The Fcho2 coding sequence (Dharmacon clone ID 6830607) was subcloned into the pGEX-6P-1 vector (GE Healthcare) using EcoRI and NotI restriction sites. DNA constructs for AP2βHE were assembled according to Kelly et al^18^., while those for Clathrin Heavy and Light Chains were produced using previously published methods. The pGEX-4T2 vector containing rat Epsin1 was a gift from H. McMahon^26^. Eps15ΔCC plasmid was generated by introducing SalI restriction sites to excise residues 315–480 of Eps15.The full-length human Hrb construct was cloned into a modified pGEX expression vector to generate an N-terminal GST fusion and a C-terminal His6 tag (PGEX-HrbFL-His6). The construct encodes GST followed by a thrombin cleavage site upstream of the Hrb coding sequence, with a hexahistidine tag fused at the C-terminus to allow sequential glutathione and Ni2+ affinity purification. All plasmids utilized in this work are available upon request.

### Protein Purification

Eps15 and Fcho2 were purified following the procedures described by Day et al^5^. In brief, Eps15 was produced as an N-terminal 6×His-tagged construct in E. coli BL21 (DE3) cells, while Fcho2 was expressed as an N-terminal glutathione S-transferase (GST) fusion protein in E. coli BL21 Star (DE3) pLysS cells. Bacterial cultures were grown in 2×YT medium at 30 °C for 4 hours until the optical density at 600 nm reached 0.6. Cultures were then chilled for 1 hour prior to induction with 1 mM IPTG. Expression was carried out under distinct temperature conditions: Eps15 at 12 °C for 20 hours and Fcho2 at 30 °C for 6 hours. Cells were subsequently collected by centrifugation, and cell disruption was achieved by homogenization followed by probe sonication in the respective lysis buffers.

Clathrin Heavy Chain (CHC) and Clathrin Light Chain (CLC) were purified following previously established methods^5^. In brief, *E. coli* BL21 competent cells (NEB) were transformed with the CHC expression plasmid pET28A(+)-6His-ratCHC-FL and cultured in 2×TY medium supplemented with 10×M9 salts and kanamycin (pH 7.4). Cultures were grown at 30 °C until the optical density at 600 nm reached approximately 1.4, then cooled to 12 °C and induced with 1 mM IPTG for 24 hours. Cell pellets were harvested, frozen, and stored at −80 °C until further use. For purification, cell pellets from 2 L of culture were thawed on ice and resuspended in 140 mL of lysis buffer (0.5 M Tris-HCl, pH 8.0; 5 mM TCEP; 1% Triton X-100) supplemented with a protease inhibitor cocktail. The suspension was then sonicated on ice, and the resulting lysate was batch-incubated with Ni–NTA agarose resin for 1 hour at 4 °C. After washing, bound proteins were eluted and concentrated via ammonium sulfate precipitation. The precipitate was dissolved and subjected to size-exclusion chromatography on a Superose 6 column. Purified CHC fractions were pooled, reconcentrated by ammonium sulfate precipitation, dialyzed, and stored as liquid nitrogen pellets at −80 °C.

Clathrin Light Chain (CLC) was co-expressed with Clathrin Heavy Chain (CHC) and purified by selective thermal denaturation of CHC. E. coli BL21 competent cells were co-transformed with the plasmids pET28A(+)-6His-ratCHC-FL and pBAT4 (untagged CLCA1). Expression and initial purification followed the same procedures as for CHC, with the culture medium additionally supplemented with 100 µg/mL ampicillin to maintain the CLC plasmid. After the first ammonium sulfate precipitation, the protein pellet was resuspended and dialyzed overnight at 4 °C against buffer containing 0.5 M Tris-HCl (pH 8.0), 1 mM EDTA, and 5 mM DTT. The dialyzed sample was subsequently heated to 90 °C for 5 minutes to denature and precipitate CHC, which was removed by centrifugation. Purified CLC was supplemented with DTT to a final concentration of 5 mM and stored as liquid nitrogen pellets at −80 °C. AP2βHE was expressed and purified as described previously with minor adjustments. In brief, AP2βHE was transformed into E. coli BL21(DE3)pLysS cells and expressed at 22 °C overnight. Cells were lysed in Tris buffer (50 mM Tris-HCl, pH 8.0; 1 M NaCl; 10% glycerol) supplemented with protease inhibitors. The resulting lysates were incubated with glutathione Sepharose resin, washed extensively with high-salt buffer, and eluted through cleavage with HRV protease to release the target proteins. Following tag removal via protease digestion, proteins were further purified by size-exclusion chromatography.

Epsin1 was purified following previously described protocols with slight modifications^33^. Briefly, proteins were expressed in E. coli cultures at 18 °C overnight and purified from bacterial lysates by binding to glutathione–Sepharose resin. The resin was washed extensively with buffer containing 25 mM HEPES (pH 7.4), 150–300 mM NaCl, 2 mM EDTA, and 2 mM dithiothreitol (DTT), followed by equilibration in a final buffer of 25 mM HEPES (pH 7.4), 150 mM NaCl, 2 mM EDTA, and 2 mM DTT. Bound proteins were cleaved from the resin using thrombin by incubation for 2 hours at room temperature and subsequently overnight at 4 °C. Thrombin was removed through adsorption with benzamidine–Sepharose (GE Healthcare) for 30 minutes at room temperature. The eluted proteins were concentrated and dialyzed for 2 hours at room temperature and then overnight in buffer containing 25 mM HEPES (pH 7.4), 150 mM NaCl, 1 mM EDTA, and 1 mM β-mercaptoethanol.

AP2βHE construct was expressed and purified following established protocols^18^. In brief, constructs were transformed into BL21(DE3)pLysS cells and induced overnight at 22 °C. Cells were lysed in 50 mM Tris pH 8.0, 1 M NaCl, 10% glycerol buffer with protease inhibitors. Lysates were loaded onto glutathione Sepharose, washed with a high-salt buffer, and eluted by on-column HRV protease cleavage. Cleaved proteins were further purified via size-exclusion chromatography For Hrb the PGEX-HrbFL-His6 construct was transformed into BL21(DE3)pLysS cells and expression was induced at 22 °C overnight. Cells were harvested and resuspended in lysis buffer (10 mM Tris pH 7.4, 200 mM NaCl) supplemented with protease inhibitors, then lysed by high pressure disruption. Clarified lysate was incubated with glutathione Sepharose resin, washed extensively in 50 column volumes of the same buffer, and bound PGEX-HrbFL-His6 was eluted by on-column thrombin cleavage at room temperature for 3 h. The thrombin-cleaved material, retaining the C-terminal His6 tag, was applied to Ni-NTA resin, washed with lysis buffer containing 20mM imidazole, and eluted with buffer containing 300mM imidazole. Eluted protein was concentrated and further purified by size-exclusion chromatography.

### Protein Labeling

Eps15, Fcho2, AP2βHE, and Epsin1, and Hrb were fluorescently labeled using amine-reactive NHS ester dyes (ATTO-Tec) in phosphate-buffered saline supplemented with 10 mM sodium bicarbonate (pH 8.3). ATTO-488 NHS ester was used for Eps15, whereas ATTO-594 NHS ester was employed for Fcho2, Hrb, AP2βHE, and Epsin1. To obtain a labeling stoichiometry of approximately 1 dye per protein molecule. Labeling reactions were carried out on ice for 15 minutes, after which proteins were buffer-exchanged into 20 mM Tris-HCl, 150 mM NaCl, and 5 mM TCEP (pH 7.5). Unreacted dye was removed using 10 kDa MWCO Amicon centrifugal filters followed by cleanup on Zeba Spin Dye and Biotin Removal columns. Before each experiment, labeled proteins were clarified by centrifugation at 100,000 × g for 10 minutes at 4 °C to eliminate aggregates. For clathrin labeling, light and heavy chains were thawed on ice and combined at a 1:1 stoichiometric ratio to generate a 3 µM solution of clathrin triskelia, which was then exchanged into 100 mM sodium bicarbonate and 20 mM β-mercaptoethanol (pH 8.2) using a Centri-Spin size-exclusion column. Alexa Fluor 647 NHS ester was added at a molar ratio of 6 dyes per triskelion, and the reaction proceeded for 20 minutes at room temperature before immediate exchange into storage buffer (10 mM Tris-HCl, 20 mM β-mercaptoethanol, pH 8.0). Free dye was removed by overnight dialysis at 4 °C using Slide-A-Lyzer MINI Dialysis devices (7 kDa MWCO) followed by passage through Zeba Spin Dye and Biotin Removal columns. Labeled clathrin was cleared by centrifugation at 100,000 × g for 10 minutes, yielding a final protein concentration of 2 µM with a labeling degree of 0.5–1 dye per triskelion. Protein and dye concentrations, as well as labeling ratios, were determined by UV–visible spectroscopy.

### Endocytic condensate formation

All condensates were assembled in buffer composed of 20 mM Tris-HCl, 150 mM NaCl, 5 mM TCEP, 1 mM EDTA, 1 mM EGTA, and 3% (w/v) PEG8000 (pH 7.5), unless otherwise indicated. For condensate assays containing only Eps15, a protein concentration of 20 µM was used. Experiments with accessory proteins utilized 20 µM Eps15 and 10 µM of the indicated accessory proteins unless otherwise stated. Sample wells were formed by adhering silicone gaskets (Grace Bio-Labs) to ultra-clean coverslips. Coverslip surfaces were passivated with PLL-PEG, followed by 10 sequential washes with 50 µL of buffer to remove unbound PLL-PEG prior to sample addition.

### Confocal microscopy

Samples for microscopy were mounted in chambers assembled using 1.6-mm-thick silicone gaskets (Grace Biolabs) on no. 1.5 glass coverslips (VWR) that had been cleaned with Hellmanex III (Hellma). Coverslip surfaces were passivated with poly-L-lysine–grafted poly(ethylene glycol) (PLL-PEG) to minimize nonspecific adhesion. Chambers were sealed with an additional small coverslip atop the gasket to prevent evaporation during imaging. Fluorescence imaging was performed on an Olympus SpinSR10 spinning-disk confocal microscope equipped with a Hamamatsu Orca Flash 4.0V3 sCMOS camera. Fluorescence recovery after photobleaching (FRAP) was executed using the integrated Olympus FRAP unit with a 405-nm laser. Data analysis was conducted in ImageJ, where recovery curves were normalized to pre-bleach maximum intensity and fitted to quantify half-time recovery for condensates of comparable size and photobleached region. PLL–PEG was synthesized following established protocols^5^ and applied to passivate coverslips. Amine-reactive mPEG succinimidyl valerate was conjugated to poly-L-lysine at a 5:1 molar ratio of PEG to PLL. The reaction proceeded overnight at room temperature in 50 mM sodium tetraborate buffer (pH 8.5), after which the product was buffer exchanged into PBS (pH 7.4) and stored at 4 °C.

Protein partitioning (Kp) values within condensates were determined using the following equation:

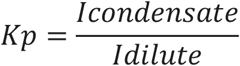

Where Icondensate is the measured fluorescence intensity within the condensate phase and Idilute is the measured fluorescence intensity within the surrounding dilute phase. Fluorescence intensity measurements were assumed to be directly proportional to local protein concentration in both the condensate and dilute phases following subtraction of the contribution from camera noise, enabling the intensity ratio to serve as a reliable proxy for the true concentration partition coefficient. For all fluorescence measurements camera noise was subtracted from overall intensity to ensure accurate measurements. By definition, Kp is a positive number. If Kp=1, it indicates that the protein is equally distributed between condensate and dilute phases. If Kp>1, the protein is enriched inside condensates compared to the surrounding solution, indicating preferential partitioning. Conversely, Kp<1 means the protein is depleted within condensates relative to the dilute phase, reflecting exclusion. In this study, Kp serves as a relative measure of partitioning to compare network density.

### Dynamic light scattering

Particle size distributions of clathrin and accessory proteins under varying pH and buffer conditions were characterized using a Malvern Zetasizer NanoZS instrument running Zetasizer software version 7.13 (laser wavelength: 633 nm; detection angle: 173° backscatter). Measurements were performed in Brand MicroUV cuvettes containing 50 µL sample volumes.

## Acknowledgements

This work was supported by the National Institutes of Health through R35GM139531 (Stachowiak), the National Science Foundation through MCB 2529782 (Stachowiak) and the Welch Foundation through F-2257 (Stachowiak). ZK and DJO were supported by Wellcome Trust grants 220597/Z/20/Z and 227915/Z/23/Z. ZK was additionally funded by GAČR 21-16786M.

## Author contributions

B.M., E.M.L., Z.K., D.O, S.S, and J.C.S. designed experiments. B.M. and J.C.S. wrote and edited the manuscript. B.M., L.W., D.O., and S.S, performed experiments. All authors consulted on manuscript preparation and editing.

## Supplementary Materials

**Supplementary Figure 1.**
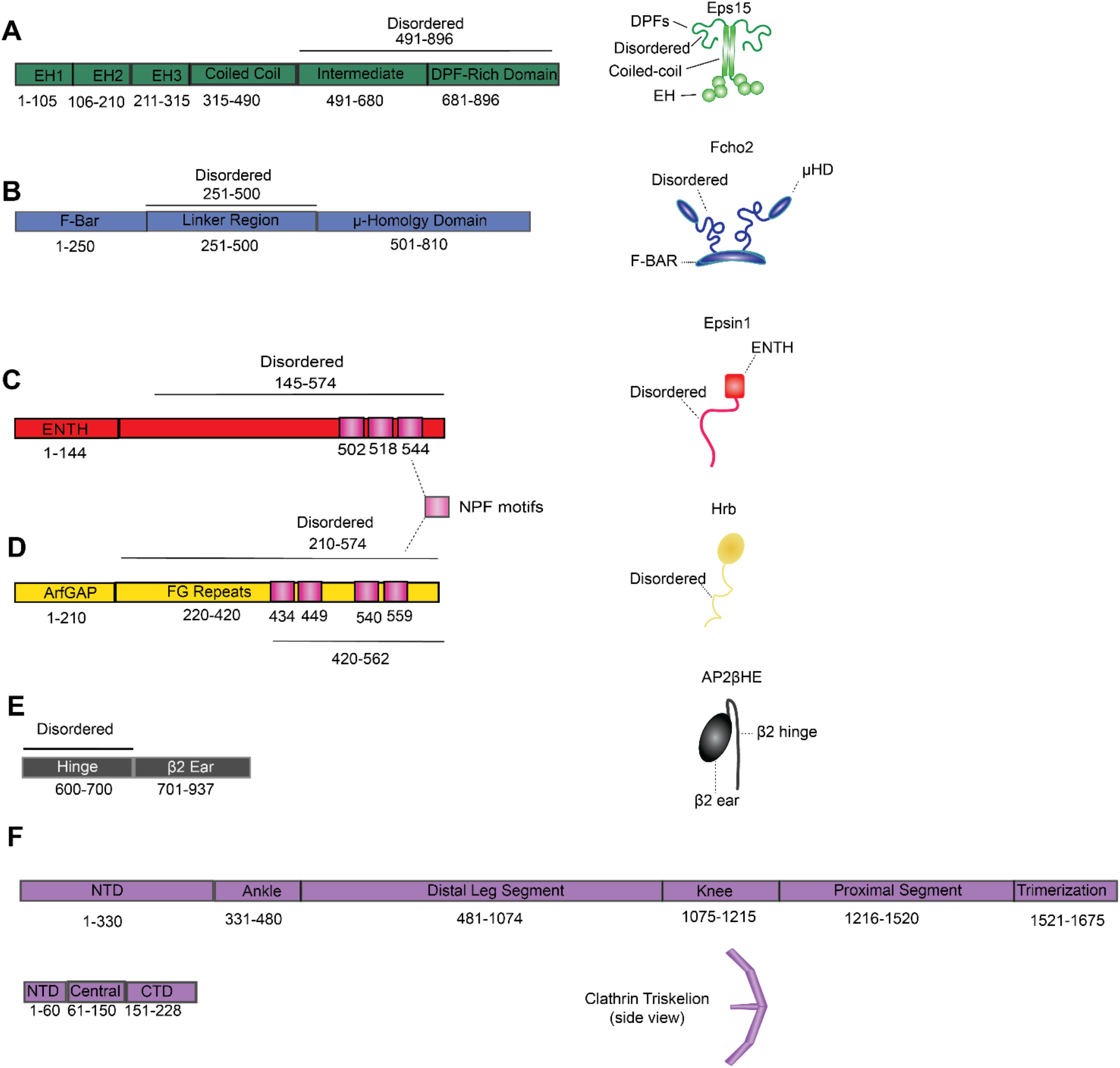
Endocytic proteins Investigated in this study. **A)** Schematic of Eps15 domain organization. **B)** Schematic of Fcho2 domain organization. **C)** Schematic of Epsin1 domain organization. **D)** Schematic of Hrb domain organization. **E)** Schematic of AP2βHE domain organization. **F)** Schematics of Clathrin Light Chain and Clathrin Heavy Chain domain organization.

**Supplementary Figure 2.**
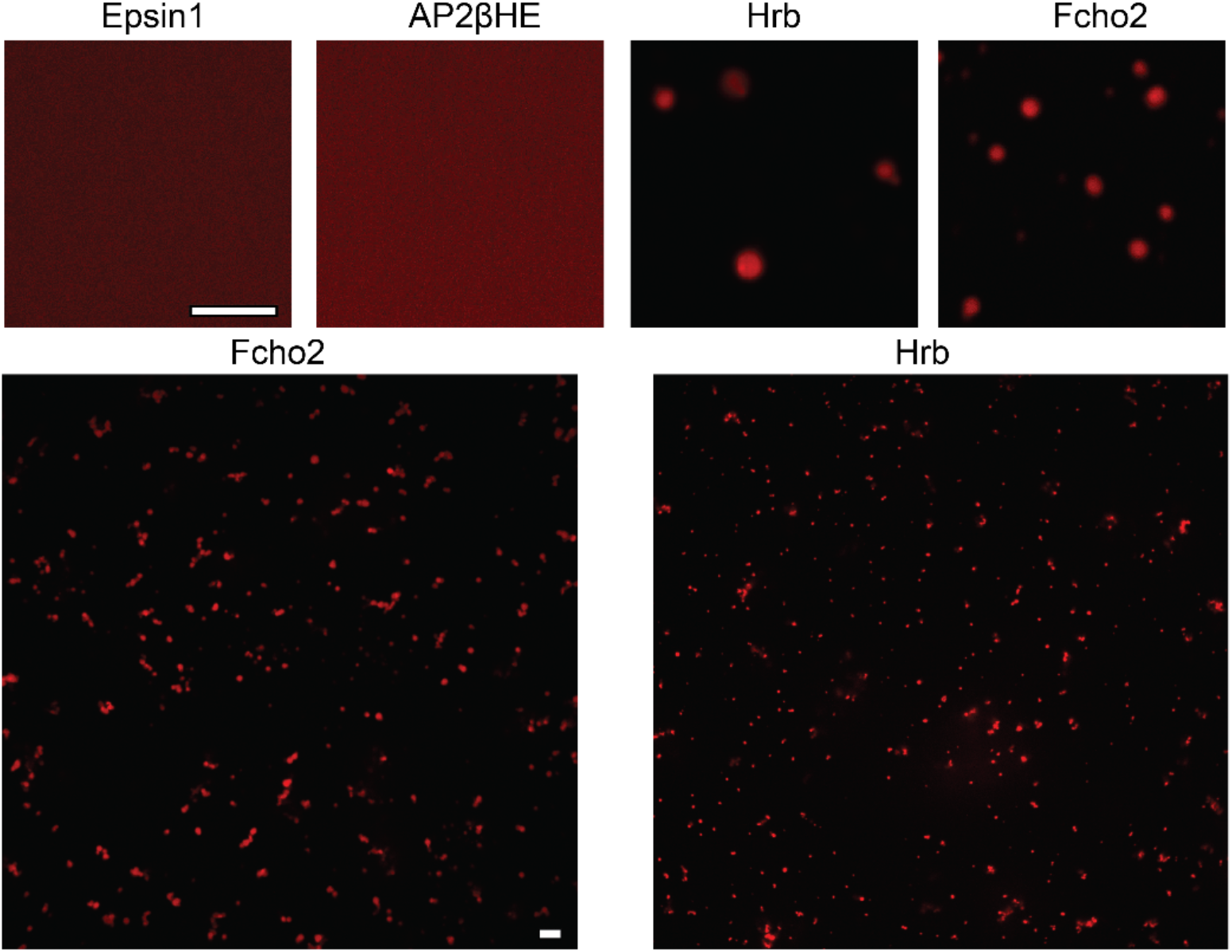
Self-interaction potential for the accessory proteins investigated in this study. Confocal microscopy images of 20 μM of the indicated accessory proteins in the presence of 3% PEG in TNEET buffer (pH 7.5 20 mM Tris-HCl, 150 mM NaCl, 1 mM EDTA, 1 mM EGTA, 5mM TCEP). For Hrb and Fcho2 a larger field of view is included to show the irregular structures.

**Supplementary Figure 3.**
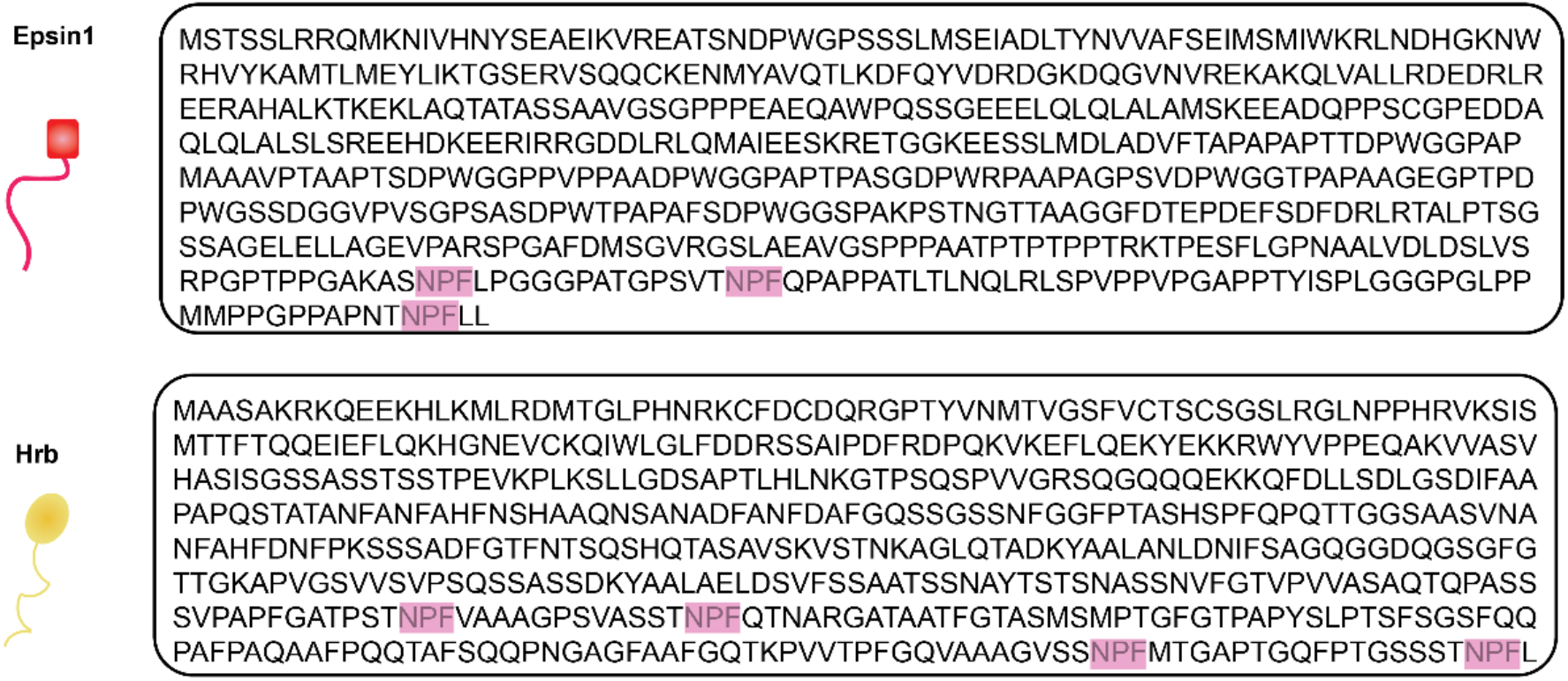
Comparison of NPF motif spacing in Epsin1 and Hrb. Amino acid Sequences for Epsin1 and Hrb highlighting the varying patterning of NPF motifs throughout the sequences.

**Supplementary Figure 4.**
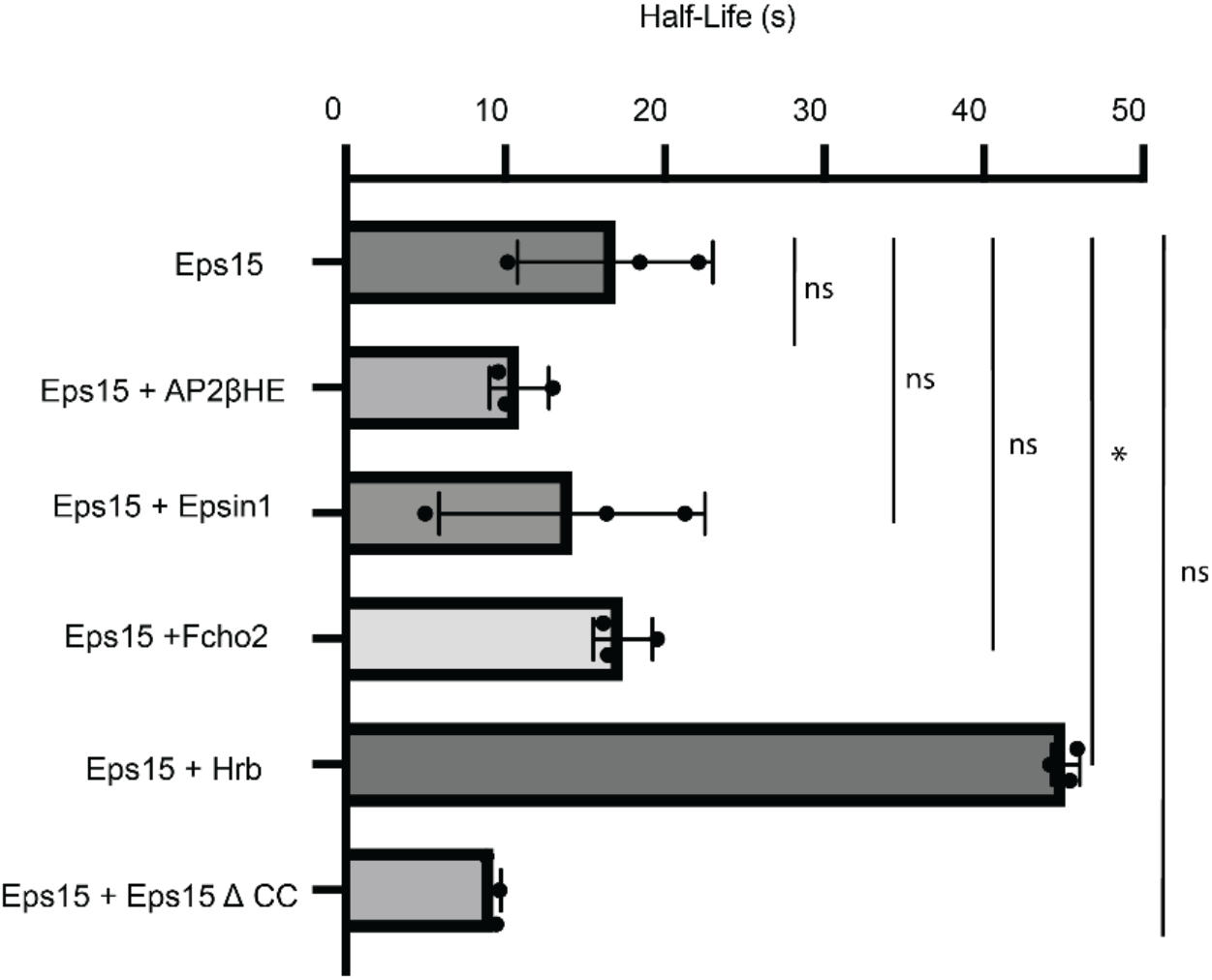
Half-Lives from FRAP experiments on condensates with varying compositions. Bar chart showing average half-lives of various condensate compositions across n=3 replicates. All condensates were formed in TNEET buffer (pH 7.5 20 mM Tris-HCl, 150 mM NaCl, 1 mM EDTA, 1 mM EGTA, 5mM TCEP) with 20 μM Eps15 and 10 μM of accessory proteins when present. GraphPad Prism Version 10.3.0 was used for graphing and curve fitting using a single phase-exponential fit to extract half-life and check significance. For statistics Welch’s two-tailed t-test was used: ns denotes no significance, * denotes *p* ≤ 0.05, ** denotes *p* ≤ 0.01, *** denotes *p* ≤ 0.001 and **** denotes *p* ≤ 0.0001. GraphPad Prism Version 10.3.0 was used for statistical analysis.

## References

(1) Pearse, B. M. Clathrin: A Unique Protein Associated with Intracellular Transfer of Membrane by Coated Vesicles. Proc Natl Acad Sci U S A 1976, 73 (4), 1255–1259. 10.1073/pnas.73.4.1255.

(2) Kaksonen, M.; Roux, A. Mechanisms of Clathrin-Mediated Endocytosis. Nat Rev Mol Cell Biol 2018, 19 (5), 313–326. 10.1038/nrm.2017.132.

(3) McMahon, H. T.; Boucrot, E. Molecular Mechanism and Physiological Functions of Clathrin-Mediated Endocytosis. Nat Rev Mol Cell Biol 2011, 12 (8), 517–533. 10.1038/nrm3151.

(4) Malady, B. T.; Papagiannoula, A.; Kamatar, A.; Sarkar, S.; Lebrun, G. T.; Wang, L.; Hayden, C. C.; Lafer, E. M.; Owen, D. J.; Milles, S.; Stachowiak, J. C. Of Condensates and Coats -Reciprocal Regulation of Clathrin Assembly and the Growth of Protein Networks. Nat Commun 2025, 16 (1), 9139. 10.1038/s41467-025-64816-x.

(5) Day, K. J.; Kago, G.; Wang, L.; Richter, J. B.; Hayden, C. C.; Lafer, E. M.; Stachowiak, J. C. Liquid-like Protein Interactions Catalyse Assembly of Endocytic Vesicles. Nat Cell Biol 2021, 23 (4), 366–376. 10.1038/s41556-021-00646-5.

(6) Kozak, M.; Kaksonen, M. Condensation of Ede1 Promotes the Initiation of Endocytosis. eLife 2022, 11, e72865. 10.7554/eLife.72865.

(7) Dragwidge, J. M.; Wang, Y.; Brocard, L.; De Meyer, A.; Hudeček, R.; Eeckhout, D.; Grones, P.; Buridan, M.; Chambaud, C.; Pejchar, P.; Potocký, M.; Winkler, J.; Vandorpe, M.; Serre, N.; Fendrych, M.; Bernard, A.; De Jaeger, G.; Pleskot, R.; Fang, X.; Van Damme, D. Biomolecular Condensation Orchestrates Clathrin-Mediated Endocytosis in Plants. Nat Cell Biol 2024, 26 (3), 438–449. 10.1038/s41556-024-01354-6.

(8) Sarkar, S.; Liu, H.-Y.; Yuan, F.; Malady, B. T.; Wang, L.; Perez, J.; Lafer, E. M.; Huibregtse, J. M.; Stachowiak, J. C. Epsin1 Enforces a Condensation-Dependent Checkpoint for Ubiquitylated Cargo during Clathrin-Mediated Endocytosis. bioRxiv February 16, 2025, p 2025.02.12.637885. 10.1101/2025.02.12.637885.

(9) Dai, Y.; You, L.; Chilkoti, A. Engineering Synthetic Biomolecular Condensates. Nat Rev Bioeng 2023, 1–15. 10.1038/s44222-023-00052-6.

(10) Ma, L.; Umasankar, P. K.; Wrobel, A. G.; Lymar, A.; McCoy, A. J.; Holkar, S. S.; Jha, A.; Pradhan-Sundd, T.; Watkins, S. C.; Owen, D. J.; Traub, L. M. Transient Fcho1/2⋅Eps15/R⋅AP-2 Nanoclusters Prime the AP-2 Clathrin Adaptor for Cargo Binding. Dev Cell 2016, 37 (5), 428–443. 10.1016/j.devcel.2016.05.003.

(11) Fazioli, F.; Minichiello, Liliana; Maťoškovā, Brona; Wong, William T.; and Di Fiore, P. P. Eps15, A Novel Tyrosine Kinase Substrate, Exhibits Transforming Activity. Molecular and Cellular Biology 1993, 13 (9), 5814–5828. 10.1128/mcb.13.9.5814-5828.1993.

(12) Parachoniak, C. A.; Park, M. Distinct Recruitment of Eps15 via Its Coiled-Coil Domain Is Required For Efficient Down-Regulation of the Met Receptor Tyrosine Kinase. J Biol Chem 2009, 284 (13), 8382–8394. 10.1074/jbc.M807607200.

(13) Cupers, P.; ter Haar, E.; Boll, W.; Kirchhausen, T. Parallel Dimers and Anti-Parallel Tetramers Formed by Epidermal Growth Factor Receptor Pathway Substrate Clone 15. J Biol Chem 1997, 272 (52), 33430–33434. 10.1074/jbc.272.52.33430.

(14) Lafer, E. M. Clathrin–Protein Interactions. Traffic 2002, 3 (8), 513–520. 10.1034/j.1600-0854.2002.30801.x.

(15) Papagiannoula, A.; Vedel, I. M.; Motzny, K.; Tengo, M.; Saiti, A.; Milles, S. Promiscuous and Multivalent Interactions between Eps15 and Partner Protein Dab2 Generate a Complex Interaction Network. bioRxiv February 25, 2025, p 2024.09.30.615448. 10.1101/2024.09.30.615448.

(16) Annunziata, O.; Asherie, N.; Lomakin, A.; Pande, J.; Ogun, O.; Benedek, G. B. Effect of Polyethylene Glycol on the Liquid–Liquid Phase Transition in Aqueous Protein Solutions. Proceedings of the National Academy of Sciences 2002, 99 (22), 14165–14170. 10.1073/pnas.212507199.

(17) Henne, W. M.; Boucrot, E.; Meinecke, M.; Evergren, E.; Vallis, Y.; Mittal, R.; McMahon, H. T. FCHo Proteins Are Nucleators of Clathrin-Mediated Endocytosis. Science 2010, 328 (5983), 1281–1284. 10.1126/science.1188462.

(18) Kelly, B. T.; Graham, S. C.; Liska, N.; Dannhauser, P. N.; Höning, S.; Ungewickell, E. J.; Owen, D. J. AP2 Controls Clathrin Polymerization with a Membrane-Activated Switch. Science 2014, 345 (6195), 459–463. 10.1126/science.1254836.

(19) Henne, W. M.; Kent, H. M.; Ford, M. G. J.; Hegde, B. G.; Daumke, O.; Butler, P. J. G.; Mittal, R.; Langen, R.; Evans, P. R.; McMahon, H. T. Structure and Analysis of FCHo2 F-BAR Domain: A Dimerizing and Membrane Recruitment Module That Effects Membrane Curvature. Structure 2007, 15 (7), 839–852. 10.1016/j.str.2007.05.002.

(20) Doria, M.; Salcini, A. E.; Colombo, E.; Parslow, T. G.; Pelicci, P. G.; Di Fiore, P. P. The Eps15 Homology (EH) Domain-Based Interaction between Eps15 and Hrb Connects the Molecular Machinery of Endocytosis to That of Nucleocytosolic Transport. J Cell Biol 1999, 147 (7), 1379–1384. 10.1083/jcb.147.7.1379.

(21) Pryor, P. R.; Jackson, L.; Gray, S. R.; Edeling, M. A.; Thompson, A.; Sanderson, C. M.; Evans, P. R.; Owen, D. J.; Luzio, J. P. Molecular Basis for the Sorting of the SNARE VAMP7 into Endocytic Clathrin-Coated Vesicles by the ArfGAP Hrb. Cell 2008, 134 (5), 817–827. 10.1016/j.cell.2008.07.023.

(22) Chaineau, M.; Danglot, L.; Proux-Gillardeaux, V.; Galli, T. Role of HRB in Clathrin-Dependent Endocytosis. J Biol Chem 2008, 283 (49), 34365–34373. 10.1074/jbc.M804587200.

(23) Chen, H.; Fre, S.; Slepnev, V. I.; Capua, M. R.; Takei, K.; Butler, M. H.; Di Fiore, P. P.; De Camilli, P. Epsin Is an EH-Domain-Binding Protein Implicated in Clathrin-Mediated Endocytosis. Nature 1998, 394 (6695), 793–797. 10.1038/29555.

(24) Snead, W. T.; Hayden, C. C.; Gadok, A. K.; Zhao, C.; Lafer, E. M.; Rangamani, P.; Stachowiak, J. C. Membrane Fission by Protein Crowding. Proceedings of the National Academy of Sciences 2017, 114 (16), E3258–E3267. 10.1073/pnas.1616199114.

(25) Drake, M. T.; Downs, M. A.; Traub, L. M. Epsin Binds to Clathrin by Associating Directly with the Clathrin-Terminal Domain: EVIDENCE FOR COOPERATIVE BINDING THROUGH TWO DISCRETE SITES*. Journal of Biological Chemistry 2000, 275 (9), 6479–6489. 10.1074/jbc.275.9.6479.

(26) Ford, M. G. J.; Mills, I. G.; Peter, B. J.; Vallis, Y.; Praefcke, G. J. K.; Evans, P. R.; McMahon, H. T. Curvature of Clathrin-Coated Pits Driven by Epsin. Nature 2002, 419 (6905), 361–366. 10.1038/nature01020.

(27) Busch, D. J.; Houser, J. R.; Hayden, C. C.; Sherman, M. B.; Lafer, E. M.; Stachowiak, J. C. Intrinsically Disordered Proteins Drive Membrane Curvature. Nat Commun 2015, 6 (1), 7875. 10.1038/ncomms8875.

(28) Owen, D. J.; Vallis, Y.; Pearse, B. M. F.; McMahon, H. T.; Evans, P. R. The Structure and Function of the Β2-Adaptin Appendage Domain. EMBO J 2000, 19 (16), 4216–4227. 10.1093/emboj/19.16.4216.

(29) Böcking, T.; Aguet, F.; Rapoport, I.; Banzhaf, M.; Yu, A.; Zeeh, J. C.; Kirchhausen, T. Key Interactions for Clathrin Coat Stability. Structure 2014, 22 (6), 819–829. 10.1016/j.str.2014.04.002.

(30) Alberti, S.; Gladfelter, A.; Mittag, T. Considerations and Challenges in Studying Liquid-Liquid Phase Separation and Biomolecular Condensates. Cell 2019, 176 (3), 419–434. 10.1016/j.cell.2018.12.035.

(31) Ye, W.; Lafer, E. M. Clathrin Binding and Assembly Activities of Expressed Domains of the Synapse-Specific Clathrin Assembly Protein AP-3 *. Journal of Biological Chemistry 1995, 270 (18), 10933–10939. 10.1074/jbc.270.18.10933.

(32) Stetefeld, J.; McKenna, S. A.; Patel, T. R. Dynamic Light Scattering: A Practical Guide and Applications in Biomedical Sciences. Biophys Rev 2016, 8 (4), 409–427. 10.1007/s12551-016-0218-6.

(33) Zeno, W. F.; Baul, U.; Snead, W. T.; DeGroot, A. C. M.; Wang, L.; Lafer, E. M.; Thirumalai, D.; Stachowiak, J. C. Synergy between Intrinsically Disordered Domains and Structured Proteins Amplifies Membrane Curvature Sensing. Nat Commun 2018, 9 (1), 4152. 10.1038/s41467-018-06532-3.

